# Cell-type-specific silence in thalamocortical circuits precedes hippocampal sharp-wave ripples

**DOI:** 10.1101/2021.05.05.442741

**Authors:** Anna R. Chambers, Christoffer Nerland Berge, Koen Vervaeke

**Affiliations:** Department of Physiology; Institute of Basic Medical Sciences, University of Oslo, Norway

## Abstract

Memory consolidation requires the encoding of neocortical memory traces, which is thought to occur during hippocampal oscillations called sharp-wave ripples (SWR). Evidence suggests that the hippocampus communicates memory-related neural patterns across distributed cortical circuits via its major output pathways. Here, we sought to understand how this information is processed in the retrosplenial cortex (RSC), a primary target circuit. Using patch-clamp recordings from mice during quiet wakefulness, we found that SWR-aligned synaptic modulation is widespread but weak, and that spiking responses are sparse. However, using cell type and projection-specific two-photon calcium imaging and optogenetics, we show that, starting 1-2 seconds before SWR, superficial inhibition in RSC is reduced, along with thalamocortical input. We propose that pyramidal dendrites experience a period of decreased local inhibition and subcortical interference in a seconds-long time window preceding hippocampal SWR. This may aid communication of weak and sparse SWR-aligned excitation between the hippocampus and neocortex, and promote the selective strengthening of memory-related connections.

## Introduction

According to prominent models of memory consolidation, hippocampal ensembles are repeatedly reactivated during sleep and quiet wakefulness to strengthen memory traces in neocortical circuits (McClelland et al., 1995; Wilson and McNaughton, 1994). This reactivation occurs during brief, high-frequency oscillatory events in the hippocampus known as sharp-wave ripples (SWR) (Buzsaki et al., 1992; Foster and Wilson, 2006; Nádasdy et al., 1999). While a wealth of information supports the role of SWR in memory formation, the circuit mechanisms underlying this hippocampal-neocortical dialogue remain unclear.

An important output of the hippocampus runs via the subiculum to the retrosplenial cortex (RSC) (Sugar et al., 2011; Wyss and Van Groen, 1992). Because of its connectivity with the hippocampus and other cortical areas (Vann et al., 2009), RSC could play a key role in coordinating the interaction between the hippocampus and the neocortex (Nitzan et al., 2020). Recent work shows that RSC is indeed active during SWR (Karimi Abadchi et al., 2020; Khodagholy et al., 2017) and is important for the storage and recall of contextual memories (Cowansage et al., 2014). However, based on electrophysiology studies in RSC (Alexander et al., 2018), as well as similar investigations in hippocampal area CA1 (Mizuseki and Buzsáki, 2013) and in the prefrontal and entorhinal cortices (Jadhav et al., 2016; Ólafsdóttir et al., 2016), spiking activity aligned to SWR appears to be temporally and spatially sparse. Moreover, patch-clamp recordings from CA1 pyramidal cells show that synaptic responses during SWR are brief, small amplitude, and dominated by inhibition (Gan et al., 2017; Hulse et al., 2016). Weak synaptic excitation could explain the sparseness of cortical responses, but also raises the question of how such sparse information can propagate through cortical circuits without interference. During quiet wakefulness in particular, the signal-to-noise ratio of cortical activity is relatively low (McGinley et al., 2015). Memory-related input to cortical dendrites likely competes for post-synaptic influence with top-down and bottom-up sensory information from the thalamus and other structures (Cauller, 1995; Cruikshank et al., 2012; Douglas and Martin, 2004).

Whole-brain fMRI and electrophysiological studies in monkeys and rodents, respectively, have suggested that subcortical structures, such as the thalamus, are transiently silent prior to SWR (Logothetis et al., 2012; Ramirez-Villegas et al., 2015; Yang et al., 2019). This subcortical silence could enhance SWR-related information transfer by reducing the amount and variance of non-mnemonic excitatory drive to cortical pyramidal dendrites during critical windows for memory consolidation. Modest SWR-aligned excitation could thus ‘stand out’ against a backdrop of lower and less variable input. However, because both excitatory and inhibitory neurons across multiple layers receive dense subcortical projections, the net effect of subcortical silence on the excitability of RSC circuits is difficult to predict.

In this study, we explore these questions with a combination of in-vivo and in-vitro whole-cell recordings, cell-type specific two-photon calcium imaging, optogenetics, and local field potential measurements in awake mice. We find that, in RSC, SWR-aligned synaptic modulation is widespread but weak, and often preceded by a 1-2 second hyperpolarization. RSC pyramidal neurons that are silent before SWR show a larger activation during SWR. This pre-SWR silence was particularly clear in the distal dendrites of RSC pyramidal neurons and in superficial inhibitory neurons. Using optogenetics, we attribute the pre-SWR silence at least in part to a robust silence of thalamocortical inputs. Taken together, the combination of (1) a reduction in thalamocortical excitation and (2) a reduction in local inhibition could prioritize sparse reactivation-related circuit activity and generate a conducive environment for memory related dendritic plasticity.

## Results

### Synaptic SWR responses in RSC are widespread but modest, and often preceded by a membrane potential hyperpolarization

To investigate synaptic responses in RSC during SWR, we performed whole-cell patch-clamp recordings in awake, head-restrained mice (Figure 1A, Figure S1A). We targeted neurons in all layers of the agranular (dorsal) RSC and simultaneously recorded SWR with an extracellular electrode in area CA1 of the ipsilateral hippocampus. Mice could walk and groom on a treadmill, except during whole-cell access, when the treadmill was immobilized to increase recording stability. SWR rates ranged from 0.06 to 0.39 Hz (Figure 1B, C, mean = 0.16 ± 0.02 Hz, 11 mice), similar to previously reported data in unrestrained awake rodents (Buzsáki, 2015). We observed synaptic responses during SWR in the majority of recorded RSC neurons (Figure 1D; 30 cells), which included both depolarizations (13/30 neurons) and hyperpolarizations (10/30 neurons) (Figure 1D). These SWR-aligned responses were, however, modest in amplitude (Figure S1B; range: −3.3 mV to 2.3 mV, mean hyperpolarization: 2.16 ± 0.29 mV, mean depolarization 1.66 ± 0.21 mV).

**Figure 1.**
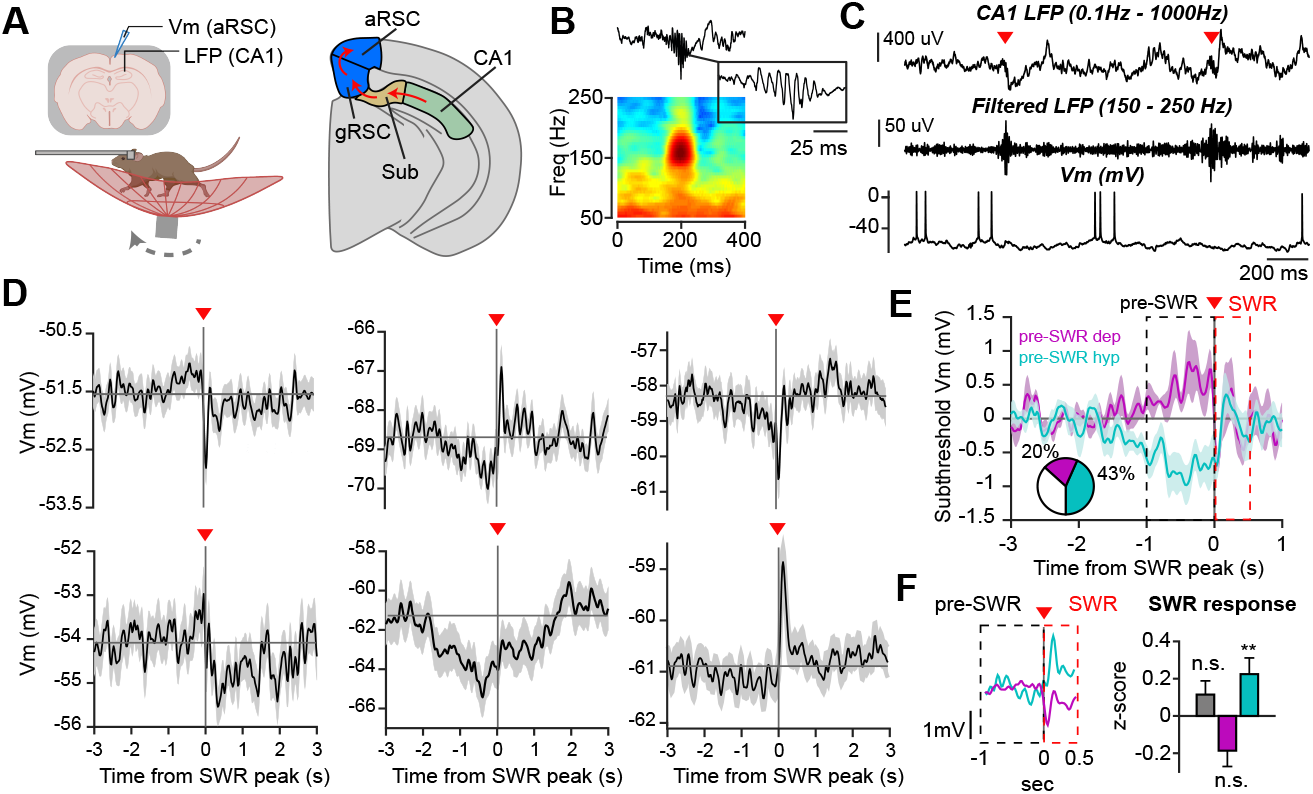
Synaptic SWR responses in RSC are widespread but modest, and often preceded by a membrane potential hyperpolarization. (**A**) Left: Experimental configuration. Simultaneous whole-cell patchclamp recording from agranular RSC (aRSC) and local field potential (LFP) recording from hippocampal area CA1, in awake head-restrained mice that can rest or walk on a treadmill. Right: Pathway connecting area CA1 with aRSC, via subiculum (Sub) and granular RSC (gRSC). (**B**) Top: Example SWR. Bottom: Average spectrogram of all SWR in one 10-minute session (n = 51 SWR). (**C**) Simultaneously recorded example traces. From top to bottom: CA1 LFP recording, filtered LFP, membrane potential (Vm) of RSC cell. Red arrowheads show SWR peak. (**D**) Average Vm ± s.e.m. of example RSC neurons aligned to SWR peak. Multiple examples are shown to illustrate the diversity of responses. (**E**) Average Vm ± s.e.m. of cells that show a significant depolarization (6/30 cells magenta, 11 mice) or hyperpolarization (13/30 cells, cyan) during the pre-SWR period. Pre-SWR and SWR periods are indicated with dashed lines. Inset shows percentage of cells in each category. (**F**) Left: Average Vm of two example cells. When the pre-SWR Vm is used as a baseline, a depolarizing ramp (magenta) is often followed by an inhibition during the SWR, while a hyperpolarizing ramp (cyan) is often followed by an excitation. Right: Average SWR response of cells without a pre-SWR change (grey, 11 cells, p = 0.17), with a pre-SWR depolarization (magenta, 6 cells, p = 0.16) or a pre-SWR hyperpolarization (cyan, 13 cells, p = 0.01). One sample Wilcoxon signed rank test).

Unexpectedly, we observed a slow, ramping response starting 1-2 seconds before the SWR in 63 % of neurons (Figure 1D, E). The majority of these pre-SWR ramps were hyperpolarizations (13/30 cells; vs 6/30 cells with depolarizing ramps; Figure 1E, inset). Since the mice were awake, we could not attribute the pre-SWR response to a toggling between two preferred membrane potentials, known as up and down states, that sometimes precedes SWR during sleep (Battaglia, 2004; Steriade et al., 2001) (see Vm traces in Figure 1C, Figure S1B). When considering the 1-second pre-SWR period as a baseline (Figure 1F, left), hyperpolarizing ramps tended to be followed by a relative depolarization during SWR (within 500 ms after SWR peak), an effect that was significant across the 13 cells with pre-SWR hyperpolarizations (Figure 1F, cyan bar; p = 0.01, one-sample Wilcoxon signed rank test). We observed an opposite trend for cells with depolarizing ramps, however this was not significant across the population. (Figure 1F, magenta bars; p = 0.16, one-sample Wilcoxon signed rank test). Due to the sparse firing of most cells, pre-SWR Vm modulations were sometimes, but not always, reflected in spiking output (Figure S1C1-4). Only 7 % of cells displayed pre-SWR decreases in spiking, in contrast to the larger proportion that displayed a pre-SWR hyperpolarization. In summary, we observed weak synaptic responses in most recorded neurons, with a substantial number of cells displaying a pre-SWR hyperpolarization coupled to a depolarization during the SWR.

### Pyramidal neurons with decreased activity prior to SWR show larger activation during SWR

Next, we investigated whether the modulation of the subthreshold membrane potential was reflected in the spiking activity of specific cell types in RSC. We used two-photon microscopy and calcium indicators to measure neural activity from a large number of identified cells (see Experimental Procedures). To target pyramidal neurons, we expressed the calcium indicator GCaMP6f under the CaMKII-promotor, or we used Thy1-GCaMP6s mice (Dana et al., 2014). We examined neurons in layer 2/3 using a glass window implanted over the agranular RSC (Figure 2A, 4 mice). In separate mice, to overcome the maximum imaging depth of two-photon microscopy (approximately 300-400 µm), we examined neurons in layer 5 through a microprism, enabling imaging up to approximately 800 µm deep (3 mice; (Andermann et al., 2013); Figure 2B). To focus our analysis on putative spiking activity with improved temporal resolution, and avoid spurious effects due to hemodynamic signal changes, we deconvolved the fluorescence signals ((Pnevmatikakis et al., 2016); Figure 2C).

**Figure 2.**
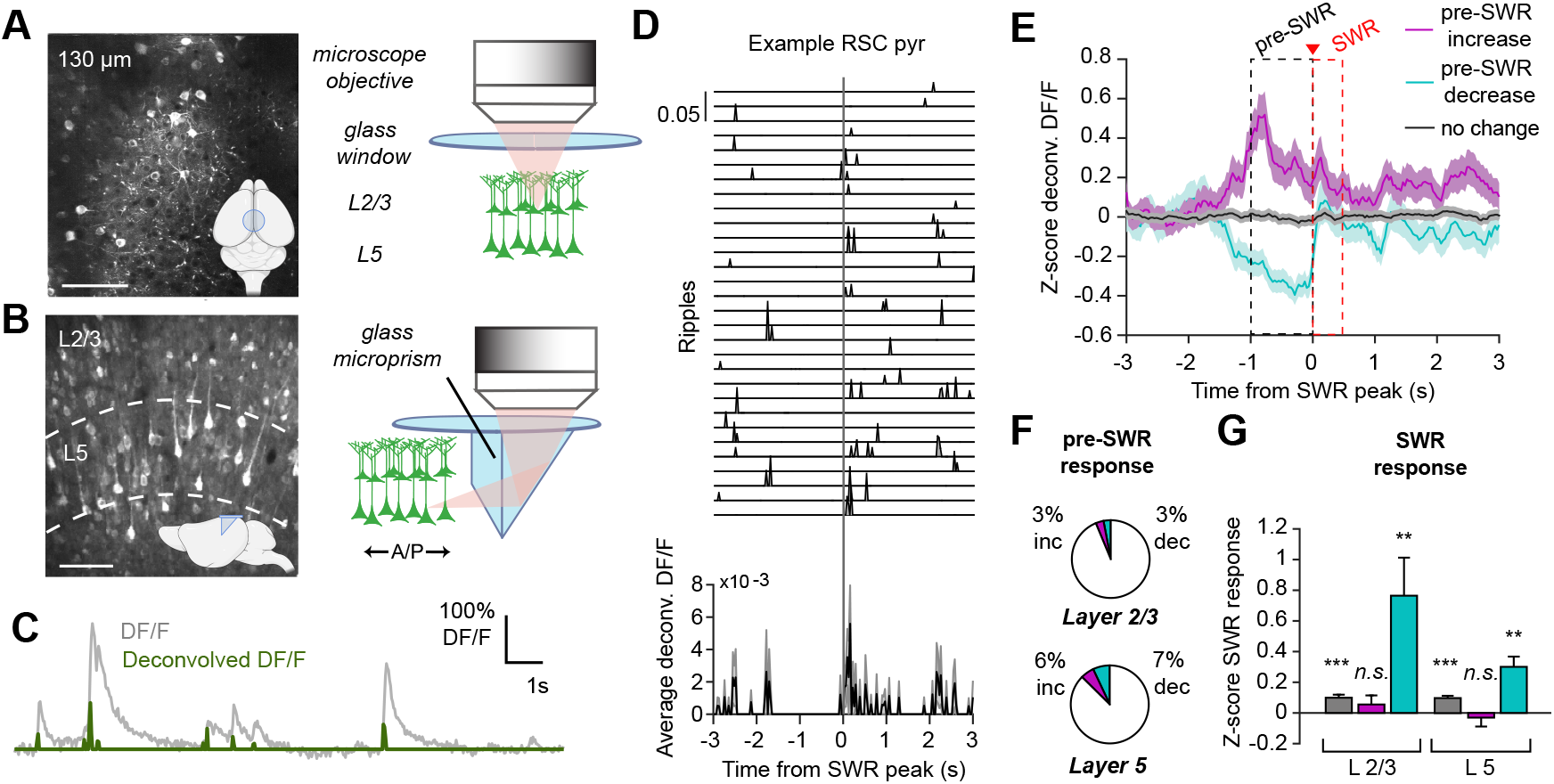
Pyramidal neurons with decreased activity prior to SWR show larger activation during SWR. (**A**) Left: Example fluorescence image of L2/3 cells in aRSC that express GCaMP6f under the CaMKII promotor (using viral delivery). Inset shows the imaging window on the mouse brain. Scale bar = 100 µm. Right: Configuration to image L2/3 cells (4 mice). (**B**) Left: Example fluorescence image of L2/3 and L5 cells in aRSC using a glass microprism (using Thy1-GCaMP6s mice). The microprism faces a coronal section of aRSC. Inset shows the location of the glass window and microprism on the mouse brain from a sagittal view. Scale bar = 100 µm. Right: Configuration to image L5 cells (3 mice). (**C**) Example fluorescence trace (fractional change in fluorescence, DF/F) from an example neuron expressing GCaMP6f under the CaMKII promotor (grey), and its deconvolved DF/F trace (green). (**D**) SWR-aligned deconvolved DF/F (top) and average +/- s.e.m. (bottom) of an example L2/3 pyramidal neuron. Note the silence before SWR, and the excitation during SWR. (**E**) Average ± s.e.m. response from all RSC pyramidal neurons (L2/3 and L5 combined) that either showed a significant decrease (cyan) or increase in fluorescence (magenta), or no change (grey) in the 1-second period before SWR. (**F**) Percentage of pyramidal neurons showing a pre-SWR modulation (L2/3 cells: 64/2127 decreased, 70/2127 increased; L5 cells: 99/1412 decreased, 79/1412 increased). (**G**) SWR response when using the pre-SWR period as a baseline. For cells with no pre-SWR change (grey bars): p = 6.46×10^−7^ (L2/3), 4.90×10^−11^ (L5). Cells with pre-SWR increase (magenta): p = 0.35 (L2/3), p = 0.6 (L5). Cells with pre-SWR decrease (cyan): p = 0.003 (L2/3), p = 2.20×10^−5^ (L5). One-sample t-test.

Mice could walk freely on the treadmill but were immobile for most of the time. Importantly, as locomotion is known to modulate the activity of multiple cell types in the cortex (McGinley et al., 2015), we only analyzed SWR events during prolonged periods of immobility (Figure S2A, see Experimental Procedures). Similar to the patch-clamp observations, we found pyramidal neurons that, on average, showed a 1-2 second reduction in activity beforeSWR, followed by an activation during SWR (Figure 2D). We then categorized cells according to whether they displayed a decrease, increase or no change in activity before SWR (Figure 2E, 2127 L2/3 cells, and 1412 L5 cells). The overall percentage of cells that decreased or increased activity before a SWR was similar in both cortical layers (Figure 2F). However, the cells that showed a decrease in activity also showed a larger response during SWR (Figure 2G). This held true for both L2/3 and L5 neurons. Neurons with no change in the pre-SWR period (Figure 2E, G) were also activated during SWR, though this increase in activity was significantly smaller than that observed in neurons with a pre-SWR silence. In summary, RSC pyramidal neurons could display a variety of changes in activity before SWR, but cells that showed decreased activity prior to SWR, also showed a larger activation during SWR.

### Pre-SWR silence is pronounced in the distal dendrites of pyramidal neurons

While the majority of neurons showed a pre-SWR modulation when measuring the membrane potential, far fewer neurons showed a pre-SWR modulation when measuring spiking activity. In addition, changes in the pre-SWR membrane potential were skewed towards hyperpolarizations (Figure 1E), while spiking increases and decreases were equally common (Figure 2F). We reasoned that, given the low basal firing rates of pyramidal neurons in the cortex—hence the large number of repetitions necessary to observe a suppressive effect— it is possible that pre-SWR spiking decreases were underestimated using two-photon microscopy. This could explain the fewer neurons showing a pre-SWR spiking modulation. But this also suggests that reductions in pre-SWR activity may be more apparent when measuring dendritic input, which has a higher level of basal activity.

To measure changes in synaptic input of pyramidal neurons, we imaged calcium signals in the tuft dendrites in layer 1 (L1) which is an important input layer for cortico-cortical and thalamo-cortical input (Van Groen and Wyss, 1992; Yamawaki et al., 2016) (Figure 3A-D). We restricted expression of GCaMP6f to RSC by injecting a small bolus of adenoassociated virus (AAV) and used the CaMKII promotor to specifically target pyramidal neurons (Figure 3A). When we measured bulk fluorescence changes in L1, to monitor many dendritic branches simultaneously, we observed that SWR were preceded by a reduction in fluorescence (Figure 3B, C). When averaging the bulk dendritic signal across mice (5 mice, 16 fields of view), fluorescence levels in the dendrites significantly decreased before a SWR, accompanied by a small but significant increase during the SWR (Figure 3D, inset; p = 0.012, one-sample Wilcoxon signed rank test). Importantly, bulk fluorescence signals may also represent non-calcium dependent changes such as brain movement or hemodynamic signal changes due to arousal. Therefore, in a subset of experiments, we also measured the calcium-independent fluorescence changes using an 810 nm laser beam (3 mice, 5 FOV; see Experimental Procedures). In these control experiments, we did not observe any significant changes in fluorescence either before or during SWR (Figure 3D). Next, to test a putative correlation between pre-SWR neural activity and arousal, we monitored the changes in pupil diameter in a subset of mice (2 mice, Figure S2B). In these data, the changes in SWR-aligned pupil diameter did not resemble the SWR modulations of neural activity (Figure S2C). Altogether, the somatic and dendritic imaging of RSC pyramidal neurons showed a pronounced silence 1-2 seconds before SWRs, which was most evident in the tuft dendrites in L1.

**Figure 3.**
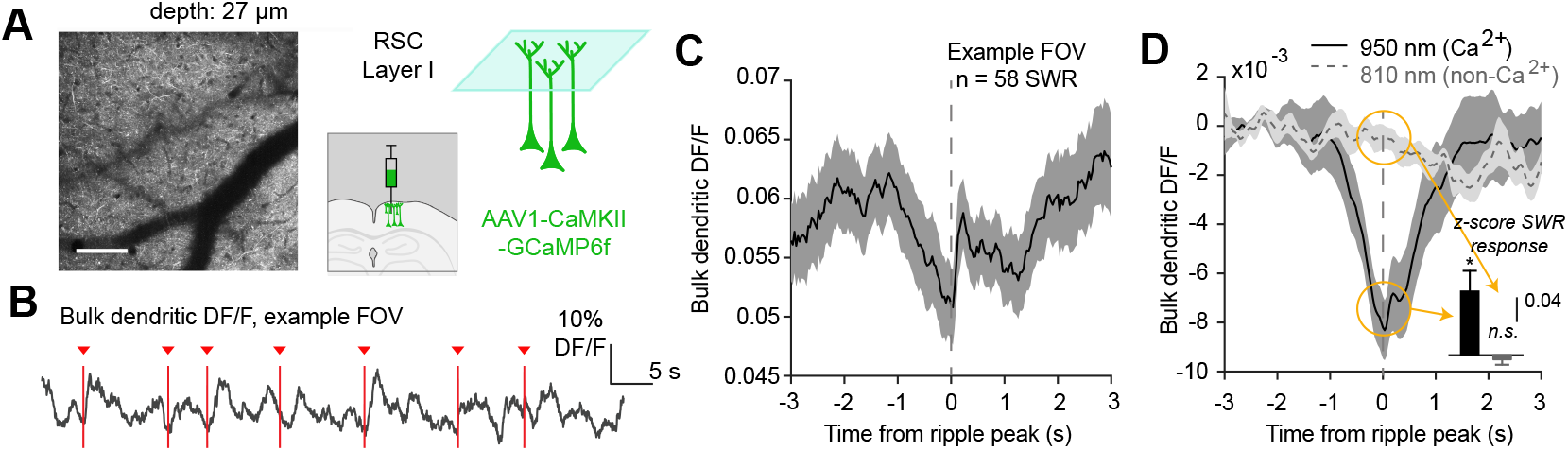
A Pre-SWR silence is pronounced in the distal dendrites of pyramidal neurons. (**A**) Left: Example fluorescence image of GCaMP6f expression in aRSC L1 (Scale bar = 100 µm). Pyramidal neurons expressed GCaMP6f under the CaMKII promotor (local viral delivery). The blue plane illustrates that L1 fields of view (FOV) contain mostly tuft dendrites from L2/3 and L5 pyramidal neurons. (**B**) Example trace of bulk DF/F (bulk = Average of all fluorescence structures in a L1 FOV). Red arrowheads indicate SWR recorded in CA1. (**C**) Average ± s.e.m. bulk DF/F aligned to SWR peak of an example FOV. (**D**) Average ±s.e.m. bulk DF/F of all FOVs (black trace; 5 mice, 16 FOV). Dashed grey line shows the Ca^2+^-independent bulk DF/F (using 810 nm excitation light: 3 mice, 5 FOV). Inset: SWR responses (time period indicated by yellow circles) for Ca^2+^-dependent and independent fluorescence measurements (black bar, p = 0.012, grey bar, p = 0.53, one-sided t-test).

### Pre-SWR modulation of inhibitory neurons is cell–type-specific and more prevalent than in excitatory neurons

The pre-SWR silence in pyramidal neuron dendrites could be due to increased inhibition or reduced excitation. To test the first possibility, we performed two-photon calcium imaging of specific inhibitory cell types using Cre mouse lines, in combination with Cre-dependent expression of GCaMP6s. We investigated L1 neurons, which are all inhibitory, and inhibitory neurons in L2/3 that express vasoactive intestinal peptide (VIP) or somatostatin (SOM), because they are key cell types that control dendritic tuft excitation (Figure 4A, B) (Tremblay et al., 2016). Further, because a subset of SOM neurons (Martinotti cells) densely innervate L1, they are of particular importance in controlling dendritic tuft excitation (Murayama et al., 2009; Wang et al., 2004). Therefore, we also imaged the axons of these cells in L1 (see Experimental Procedures).

**Figure 4.**
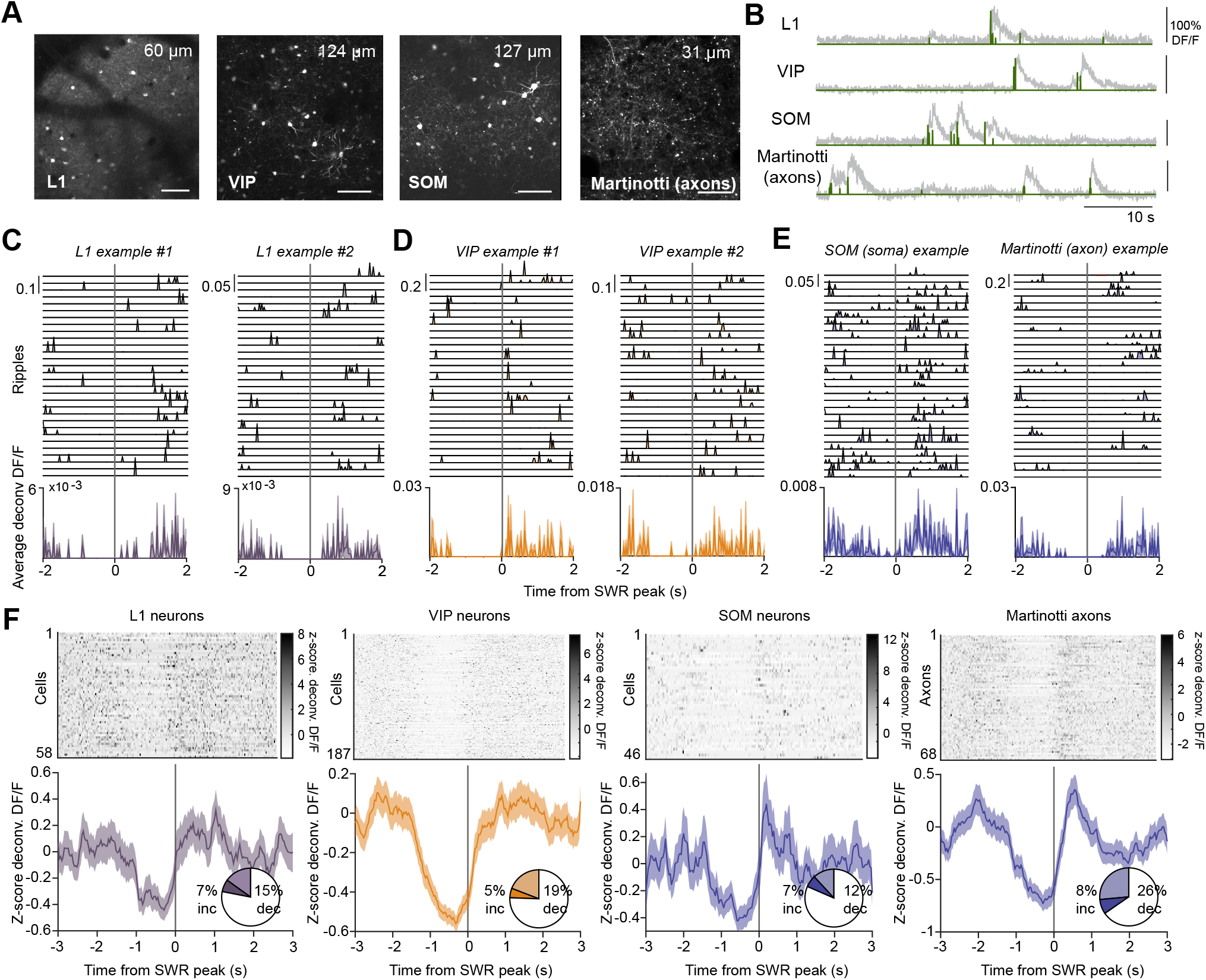
Pre-SWR modulation of inhibitory neurons is cell-type specific and more prevalent than in excitatory neurons. (**A**) Fluorescence images of GCaMP6s expression obtained by viral delivery (AAV1-CAG-flex-GCaMP6s) in Cre mouse lines using the Gad2-(for L1 neurons), VIP-, or SOM promotor. We also recorded a subset of SOM-positive neurons, called Martinotti cells, by imaging their axons in L1. Imaging depth is indicated on top. Scale bars = 100 µm. (**B**) Example fluorescence traces for each cell type (DF/F, grey) with superimposed deconvolution (green). (**C-E**) SWR-aligned deconvolved DF/F (top) and average response (bottom) of example cells showing a pre-SWR reduction in fluorescence. (**F**) Top: All cells showing a pre-SWR silence (average deconvolved DF/F). Bottom: The average ± s.e.m. response of all cells that show a pre-SWR silence. Pie charts show the percentage of cells showing a significant pre-SWR decrease or increase in fluorescence (451 L1 cells from 5 mice; 950 VIP cells from 3 mice; 363 SOM cells and 257 Martinotti axons from 3 mice).

We analyzed neural responses of the different inhibitory neurons in a 1-second time window before SWR. While we found a small percentage of cells that showed an activity increase, the majority of modulated cells showed a significant reduction in activity. We observed this across all cell types (Figure 4C-E). The percentage of cells that showed an activity increase was fairly uniform among cell types (7-8 %, Figure 4F pie charts). However, the proportion of cells showing a pre-SWR activity decrease was more cell type-dependent (Figure 4F, pie charts). A pre-SWR silence was most prominent in Martinotti axons (26 %, n = 257 axons, 3 mice) and VIP neurons (19 %; n = 950, 3 mice), followed by L1 neurons (15 %; n = 451 cells, 5 mice) and SOM neurons (12 %; n = 363 cells, 3 mice). In summary, inhibitory cells generally showed much higher proportions of SWR modulation than pyramidal neurons, with a pre-SWR silence as the dominant type of response. SOM, L1, VIP and Martinotti cells displayed varying degrees of pre-SWR silence, indicating a period of significantly reduced activity among superficial inhibitory neurons that control pyramidal cell tuft dendrites.

### Thalamocortical axons projecting to L1 show reduced activity before SWR

Since we observed a pre-SWR silence in both pyramidal neuron dendrites and in multiple types of dendrite-controlling inhibitory neurons, we hypothesized, based on recent work, that the thalamus could be a shared source of excitation that may decrease prior to SWR. (Logothetis et al., 2012; Varela and Wilson, 2020; Yang et al., 2019). To test this, we injected a retrograde AAV delivering GFP to agranular RSC and identified upstream thalamic nuclei (Figure 5A, n = 2 mice). Consistent with previous reports (Van Groen and Wyss, 1992), GFP expression was observed in cell bodies in the laterodorsal, anterodorsal, anteroventral, and anteromedial nuclei. To test whether these thalamocortical projections show a pre-SWR silence, we delivered GCaMP6s virally to the thalamus and imaged axons in L1 of RSC (Figure 5B, C, see Experimental Procedures).

**Figure 5.**
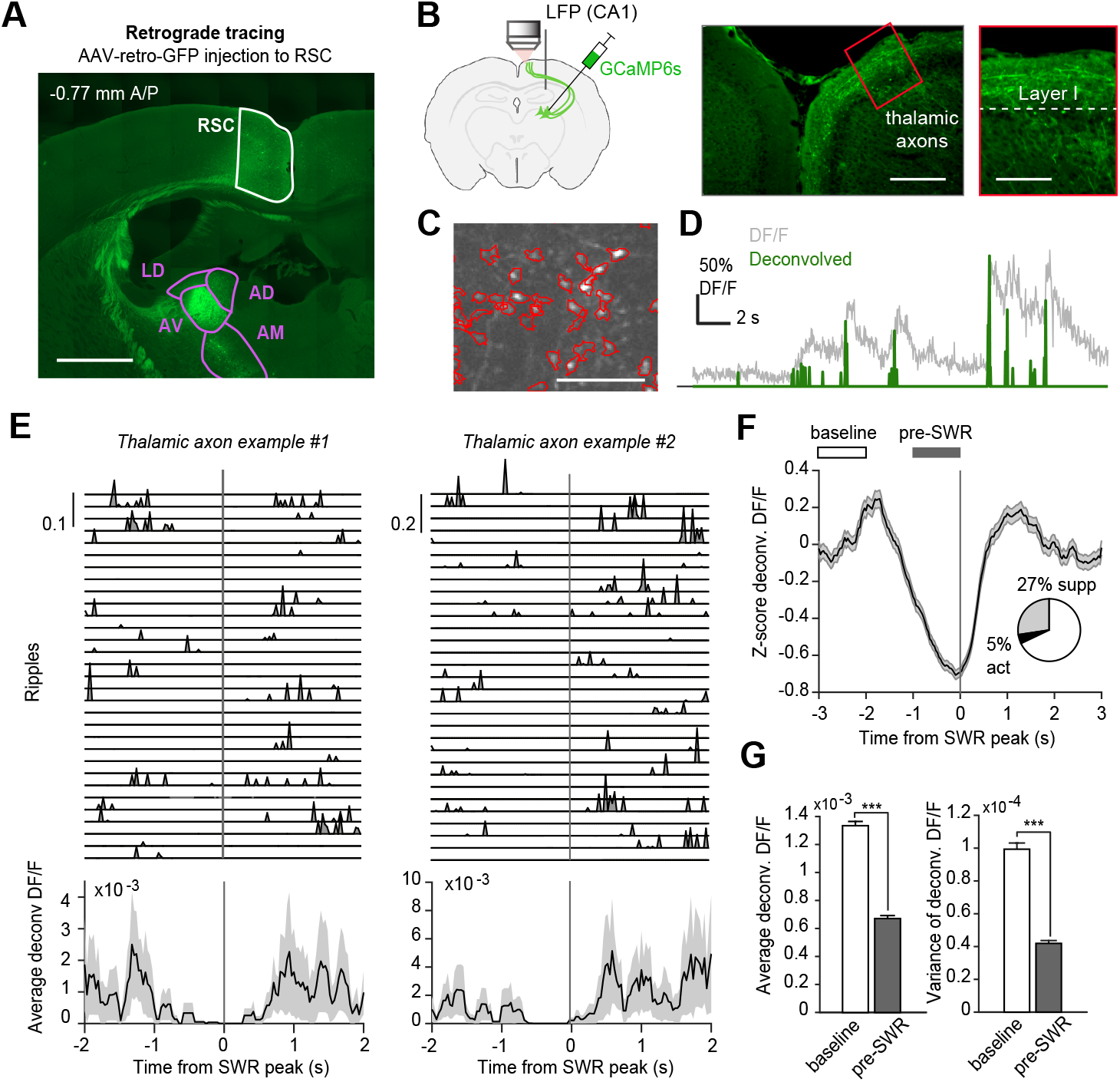
Thalamocortical axons projecting to L1 show reduced activity before SWR. (**A**) Coronal section showing an injection of rAAV2-retro-GFP in aRSC, and retrogradely-labeled GFP-positive cell bodies in thalamic nuclei (LD; laterodorsal, AV; anteroventral, AD; anterodorsal, AM; anteromedial). Scale bar = 1mm. (**B**) Left: Experimental configuration. AAV1-syn-GCaMP6s was injected in the thalamus, and axonal calcium signals were measured in L1 of aRSC with two-photon microscopy. Simultaneous LFP recordings were made in area CA1. Right: Confocal fluorescence images showing axons in aRSC L1. (**C**) Two-photon image showing thalamic axons and boutons in L1 of aRSC in an awake mouse. Scale bar = 20 µm. (**D**) Example fluorescence trace (DF/F, grey) with superimposed deconvolution (green) from a single axonal bouton recorded in an awake mouse. (**E**) SWR-aligned deconvolved responses (top), and average ± s.e.m (bottom) of example axons showing a pre-SWR reduction in fluorescence. (**F**) Average ± s.e.m. response of all axons that showed a significant pre-SWR silence. Pie chart shows the proportion of axons that show either a significant pre-SWR silence (660/2410 axons, 4 mice) or pre-SWR activation (113/2410). White and gray bars indicate time periods for calculating baseline and pre-SWR response. (**G**) Average (left) and variance (right) of the response of all axons (whether significantly modulated or not) in the 1-second period prior to SWR compared to baseline (p value (mean) = 4.30×10^−134^, p value (variance) = 2.02×10^−65^, paired t-test).

Activity in thalamocortical projections to L1 was strongly reduced in the seconds-long time window surrounding SWR (Figure 5E, F; 2364 axons, 4 mice). This reduction followed a similar time course as the inhibitory neurons and pyramidal neuron dendrites (Figures 3, 4), and was significant in 27 % of individual axons (Figure 5F, pie chart). Meanwhile, only 5 % of axons increased activity in the pre-SWR period (Figure 5F, pie chart). Across the entire axon population, we observed a pronounced decrease in both amplitude (Figure 5G, left) and variance of thalamocortical input (Figure 5G, right, p < 0.001, paired t-test). These data show that thalamocortical axons in L1 of RSC have a high basal firing rate that is markedly reduced prior to SWR.

### Thalamic silence reduces activity of L1 inhibitory neurons in RSC

The pre-SWR silence of thalamocortical input to RSC mirrors the time course of pre-SWR silence found in dendrite-controlling inhibitory neurons, but is it sufficient to cause pre-SWR silence in target cells? To test this, we first examined whether thalamocortical axons make direct, monosynaptic connections with L1 neurons. We delivered the opsin Chrimson-R (Klapoetke et al., 2014) to the thalamus so that thalamocortical axons projecting to RSC were excitable with red light (Figure 6A). We then performed patch-clamp recordings of L1 inhibitory neurons in brain slices of RSC (see Experimental Procedures). We perfused the brain slices with TTX and 4-AP to block action potentials and to enhance direct depolarization of presynaptic terminals ((Petreanu et al., 2009), Figure 6B). All (7/7) L1 neurons displayed short-latency current responses to brief pulses of light, indicating monosynaptic input from the thalamus (Figure 6C).

**Figure 6.**
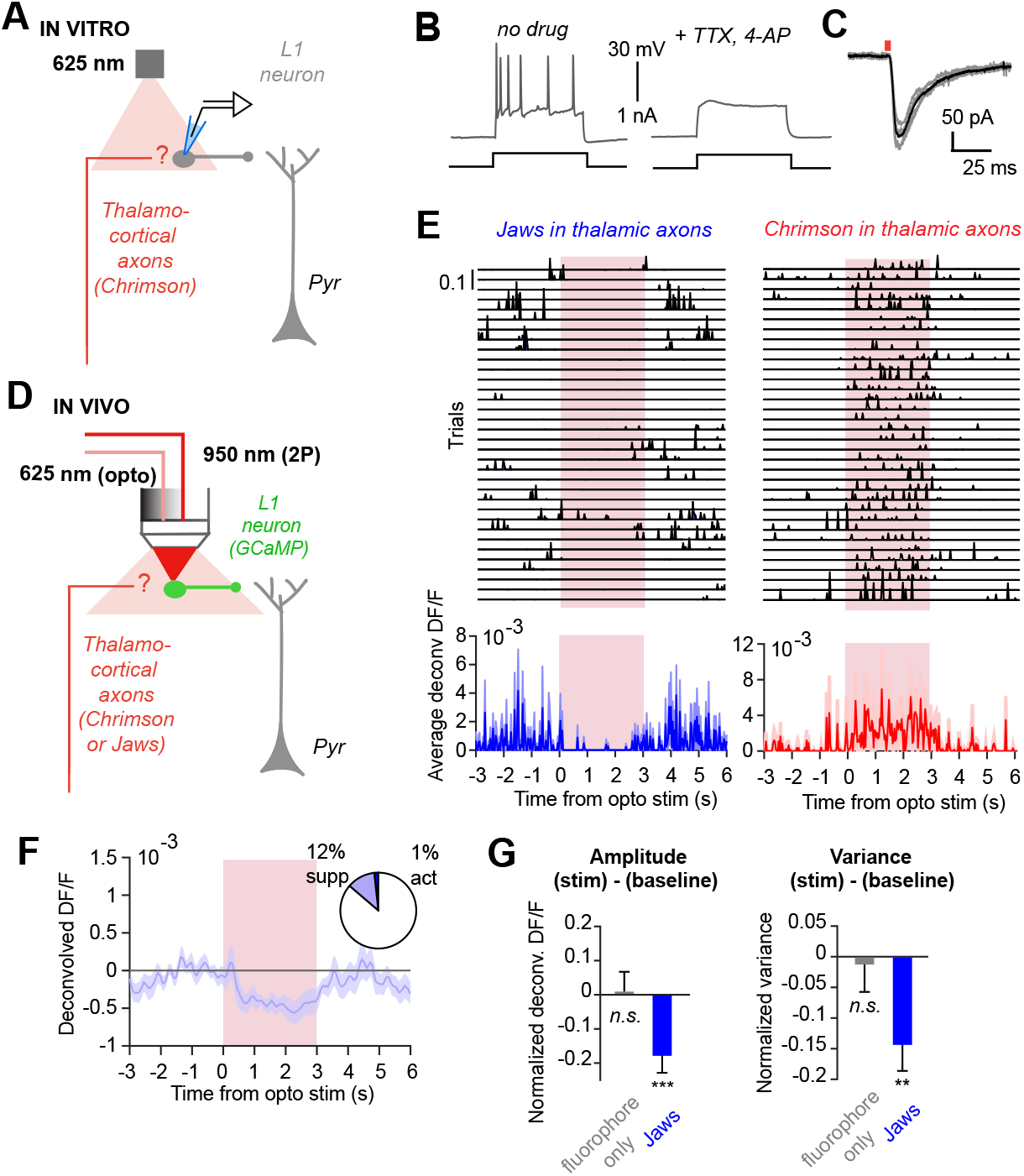
Thalamic silence reduces activity of L1 inhibitory neurons in RSC. (**A**) Experimental configuration. AAV1-syn-ChrimsonR-tdTomato was injected in the thalamus. Whole-cell patch-clamp recordings from aRSC L1 neurons were made in brain slices and red light pulses were used to excite thalamic axons. (**B**) Example Vm of an RSC L1 neuron in response to current injections before (left) and after (right) bath application of 1 µM TTX and 0.5 mM 4-AP (to block action potentials and ensure monosynaptic transmission). (**C**) Voltage clamp recording from a L1 neuron showing a typical synaptic current response to a 5 ms light pulse (black = average; grey = individual trials). All L1 neurons (7/7) showed a short latency synaptic response. (**D**) Experimental configuration for combined optogenetic modulation of thalamocortical axons and two-photon calcium imaging of aRSC L1 neurons in vivo. AAV1-syn-ChrimsonR-tdTomato or AAV5-syn-GFP-Jaws was injected in thalamus and AAV1-syn-GCaMP6s was injected in RSC. (**E**) Deconvolved DF/F responses of RSC L1 neurons during a 3-second light pulse, when thalamic axons expressed either Jaws (left example) or Chrimson-R (right example). (**F**) Average ± s.e.m. response of all L1 neurons that were suppressed during light stimulation of thalamic axons expressing Jaws (28/233 neurons, 4 mice). Pie chart shows the proportion of neurons that were either suppressed or activated during the light stimulus. (**G**) Average amplitude (left) and variance (right) of L1 neuronal activity (all cells, regardless of significant modulation), during light stimulation of thalamic axons expressing Jaws. In control experiments (181 cells, 2 mice), thalamocortical axons expressed only a fluorophore (tdTomato). (For amplitude, p value (Ctrl) = 0.88, p value (Jaws) = 0.00042; for variance, p value (Ctrl) = 0.81, p value (Jaws) = 9.58×10^−4^. One-sample t-test).

Next, we tested whether bi-directional modulation of thalamic input is mirrored by a similar modulation of L1 neurons in vivo (Figure 6D). To enable bi-directional modulation, we either expressed the excitatory opsin Chrimson-R or the in-hibitory opsin Jaws in the thalamus (Chuong et al., 2014). Using two-photon microscopy in awake mice, we monitored the activity of L1 neurons using GCaMP6s while exciting or inhibiting thalamocortical terminals with red laser light (see Experimental Procedures). Silencing thalamocortical axons with a 3-second light stimulus caused a significant suppression in 12 % of L1 neurons (5 mice, 65/538 cells, Figure 6E, F). Meanwhile, excitations were rare (1 %, 8/538 cells). This was in contrast to the activation commonly observed when thalamocortical axons expressed Chrimson-R (n = 2 mice, example in Figure 6E). Across the whole population of imaged L1 neurons, compared to control mice whose thalamocortical axons expressed only a fluorophore (tdTomato; n = 2 mice, 160 L1 neurons), the overall effect of optogenetic inhibition of thalamocortical axons was a significant reduction of the amplitude and variance of L1 neuron activity (Figure 6G, p < 0.001, one-sample t-test). Altogether, these data show that in quiet awake mice, thalamocortical input drives, at least in part, spiking in L1 neurons. Conversely, thalamic silence is sufficient to cause silence in L1 neurons.

Previous studies have suggested that silence during SWR is a widespread phenomenon observed across multiple thalamic nuclei (Logothetis et al., 2012; Yang et al., 2019). Because thalamic innervation of L1 neurons is a typical cortical feature (Cruikshank et al., 2012; Roth et al., 2016), we predicted that L1 neurons outside of RSC also display pre-SWR silence. To make large-scale measurements of L1 neuronal activity, we used AAV to express a Cre-dependent GCaMP6s in Gad2-cre mice (n = 5) and implanted cranial windows over multiple areas of dorsal cortex. We found L1 neurons with a significant pre-SWR silence in all cortical regions explored, which included posterior parietal, motor, somatosensory, and visual areas in addition to RSC (Figure S3A). Moreover, the pre-SWR silence displayed a similar time course to that seen in RSC (Figure S3B, C). In summary, the decrease in superficial inhibitory neuron activity prior to SWR observed in RSC may reflect a global cortical phenomenon related to widespread thalamic silence.

### Thalamic silence reduces calcium events in RSC pyramidal dendrites

Given the functional diversity of L1 neurons, in addition to the other excitatory and inhibitory cortical neurons that also receive thalamic input, the net effect of thalamic silence on pyramidal neuron dendrites is not straight-forward to predict. To test this, we used the opsin Jaws to inhibit thalamic axon activity in RSC while using GCaMP6 to measure dendritic tuft activity. We restricted expression of GCaMP6s to Layer 5 pyramidal neurons by injecting a retrograde virus delivering Cre in the pons (Harris and Shepherd, 2015) and a Cre-dependent GCaMP6s construct in the RSC (Figure 7A, B). This strategy offered the benefit of (1) reducing the density of labeled dendrites in L1, and thus facilitat-ing the extraction of calcium events from individual branches (Figure 7C, D); and (2) allowing us to investigate the effect of thalamocortical silence on dendritic activity in one of the major output neurons of RSC.

**Figure 7.**
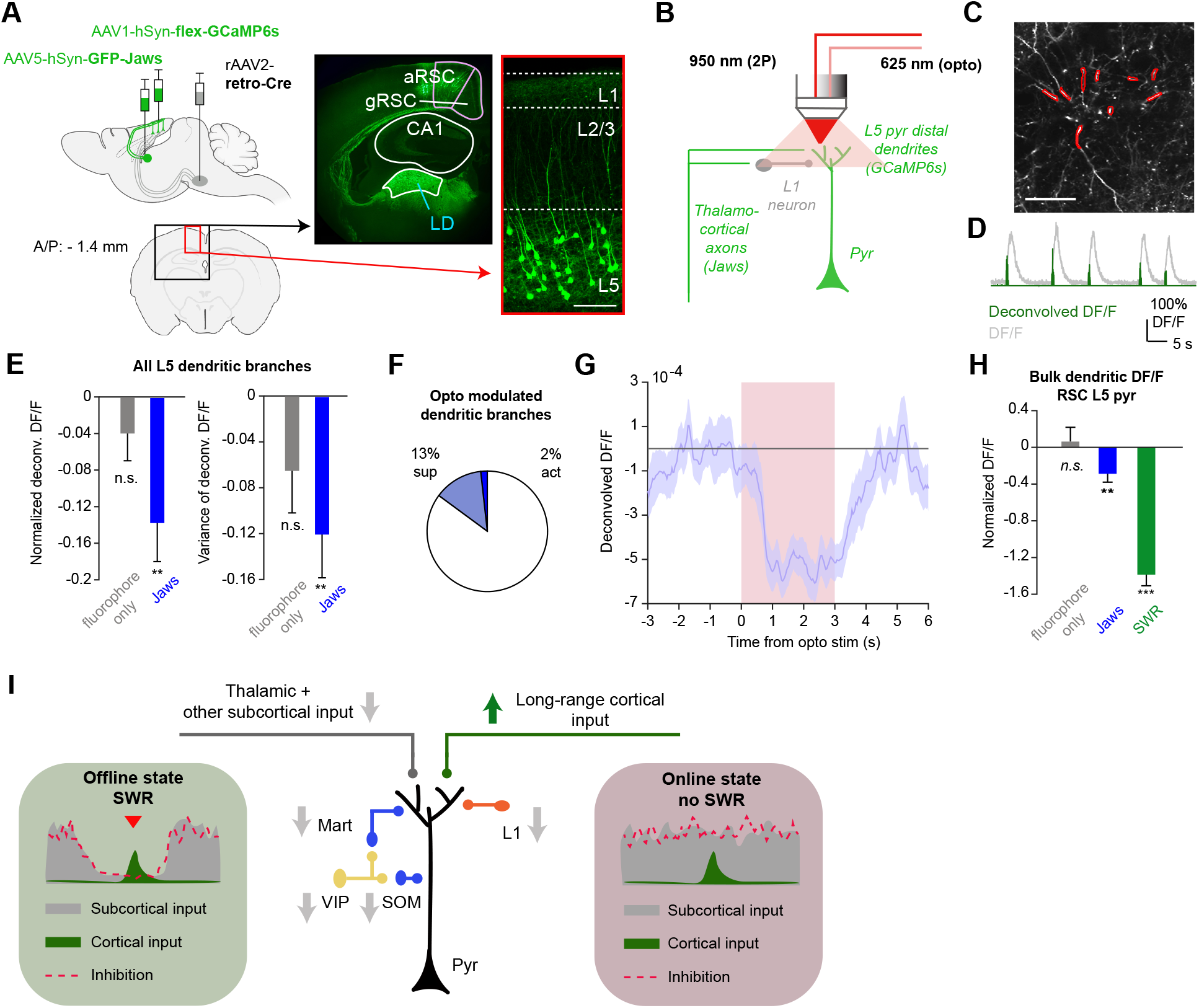
Thalamic silence reduces calcium events in RSC pyramidal dendrites. (**A**) Left: Experimental configuration for combined optogenetic inhibition of thalamocortical axons and two-photon Ca^2+^ imaging of L5 pyramidal neuron dendrites in aRSC. Right: Confocal images showing expression of GFP-Jaws in thalamus and GCaMP6s in RSC L5 neurons. Scale bar = 100 µm. (**B**) Experimental configuration for simultaneous in-vivo optogenetics and two-photon imaging experiments. Optogenetic laser stimulus was delivered through the microscope objective. (**C**) Two-photon image showing L5 pyramidal neuron tuft dendrites in RSC in an awake mouse. Dendritic branches are shown in red. Scale bar = 50 µm. (**D**) Example fluorescence trace (DF/F, grey) with superimposed deconvolution (green) from a single dendritic branch. (**E**) Average± s.e.m reduction in amplitude (left) and variance (right) of dendritic Ca^2+^ signals when inhibiting thalamic axons using Jaws (614 dendritic branches; 4 mice). In control experiments, thalamic axons expressed only a fluorophore (219 dendritic branches, 2 mice). (p value (Ctrl) = 0.22, p value (Jaws) = 0.002, One-sample t-test). (**F**) Percentage of dendritic branches that were either significantly suppressed or activated during the light stimulus. (**G**) Average ± s.e.m. deconvolved DF/F of all dendrites that were significantly suppressed during the light stimulus (red box). (**H**) Average ±s.e.m. reduction in bulk dendritic DF/F for all FOVs when inhibiting thalamocortical axons using Jaws (blue; p = 0.005), compared to a fluorophore-only (grey; tdTomato, p = 0.68), and compared to the reduction during SWR in vivo (p = 6.21×10^−15^). One-sample t-test. (**I**) RSC pyramidal neuron receiving two categories of long-range inputs at its distal dendrites: (1) thalamic + other subcortical input (left), and (2) long-range cortical input, including hippocampal input (right). Arrows indicate the activity changes in the circuit. Insets illustrate the decrease in thalamocortical input and superficial inhibition that may promote the propagation of memory-related activity in pyramidal dendrites during SWR.

We identified dendritic branches by their spiny morphology and clear calcium transients (Figure 7C, D). In response to optogenetic suppression of thalamic axon activity, a significant reduction of calcium events was observed in 13 % of individual dendritic branches (78/614 dendrites, 4 mice), while activation was rare (2 % or 11/614 dendrites, Figure 7E, F). Finally, when we considered the average response over all dendrites (Figure 7G), we found that the mean reduction in calcium activity during optogenetic Jaws stimulation was approximately 20 % of the reduction seen during SWR (p = 0.005, one-sample t-test; Figure 7H). This indicates that a silence in thalamocortical inputs could account for a sizeable portion of the pre-SWR silence in RSC dendrites in vivo.

Overall, the significant decrease in thalamocortical excitation to superficial RSC may have a dual role in promoting memory-related plasticity in pyramidal neuron dendrites (Figure 7I). First, this phenomenon results in a functional de-afferentation from thalamic input, reducing its interference with sparse cortico-cortical interactions (Figure 7I, left). Second, silence of thalamic—and possibly other subcortical—input results in decreased dendritic inhibition that would otherwise be higher if in an active, ‘online’ processing state (Figure 7I, right). Together, these effects could represent a cortical circuit mechanism for prioritizing and propagating weak excitation in order to strengthen neocortical memory traces.

## Discussion

Memory consolidation is believed to require the replay of hippocampal ensemble activity that propagates across distributed cortical circuits. We explored how excitation generated during SWR can influence cortical circuits during quiet wakefulness. We find that in RSC, which is a major hippocampal target, synaptic SWR responses are widespread but weak. However, we observed that 1-2 seconds prior to SWR, multiple cell types undergo a reduction in activity, which is most apparent in pyramidal dendrites and superficial inhibitory neurons. We attributed this, at least in part, to a transient silence in thalamocortical excitation. A reduced baseline level of activity prior to SWR in pyramidal neurons was followed by a stronger activation at SWR (Figure 1F, 2G). We propose that reduced thalamic input to cortical dendrites, coupled with lowered levels of dendritic inhibition, could aid the transfer of weak synaptic activity and the selective strengthening of memory-related connections (Figure 7I).

Previously, Logothetis et al. (2012), using fMRI in monkeys, reported widespread subcortical—particularly thala-mic—silence prior to hippocampal SWR. They proposed that subcortical silence reduces sensory interference during memory consolidation, thus increasing the signal-to-noise of hippocampal-neocortical communication. Evidence from studies using single-cell electrophysiology (Logothetis, 2015; Varela and Wilson, 2020; Wang et al., 2015; Yang et al., 2019) has supported that pre-SWR silence is not only thalamic, but a general subcortical phenomenon conserved across primates and rodents. Importantly, these silences have also been observed in neuromodulatory regions such as the median raphe nucleus (Wang et al., 2015), indicating that inputs related to arousal and movement also decrease in the pre-SWR period. Our study has expanded on this work by investigating the impact of subcortical silence on cortical circuits at the cellular and subcellular level in a cell-type-specific manner. To our knowledge, the pre-SWR silence we observed in multiple cortical cell types has not been highlighted in previous studies in awake animals (but see Battaglia (2004); Karimi Abadchi et al. (2020) during sleep/anesthesia). Previous work that used bulk imaging may have missed these changes because that method averages signals across cell types, subcellular compartments and layers. Other studies using cortical extracellular recordings (Alexander et al., 2018; Jadhav et al., 2016; Nitzan et al., 2020; Rothschild et al., 2017) have not reported a pre-SWR silence, however these recordings may have been biased toward pyramidal neurons, whose spiking activity was not strongly modulated in our experiments (Figure 2F). Finally, prior studies often focused on shorter, 100-500 ms timescales around SWR, rather than a seconds-long timescale. Nevertheless, overall, our findings broadly complement rather than contradict previous work.

Although the circuit anatomy of RSC cannot be compared directly to other neocortical regions, key similarities—such as the general distribution and function of inhibitory neurons and the patterns of superficial thalamic innervation—could allow for a generalization of our results to other cortical circuits (Figure S3). RSC L1 is a convergence site for hippocampal input (via granular RSC; Wyss and Van Groen (1992)), long-range neocortical input (from anterior cingulate, secondary motor and visual areas, and thalamic input (Van Groen and Wyss, 1992). Corticothalamic neurons typically have high basal firing rates and project densely to L1 (Figure 4A-B). Therefore, they likely provide substantial input to L1 neurons, pyramidal tuft dendrites, and possibly VIP neurons—input that decreases significantly prior to SWR. Given that these inhibitory neurons and tuft dendrites also have a high input impedance, tonic thalamic input likely raises their baseline activity levels (Figure 6E-G, Figure 7E-H). However, thalamic silence appears to have only a small impact on pyramidal cell firing, probably because pyramidal firing rates are already low (Barth and Poulet, 2012), and therefore, a subcortical silence is rather reflected by a hyper-polarization of the membrane potential (Figure 1). Similar hyperpolarizing ramps in the subthreshold membrane potential have also been reported in CA1 pyramidal neurons (Hulse et al., 2016).

The thalamus also projects to deeper layers 3 and 4 (Van Groen and Wyss, 1992). While we did not monitor the input to these layers, it likely also decreases during thalamic silence, explaining the pre-SWR activity reduction in SOM neurons (Figure 3F), and possibly PV cells, which we did not investigate. Interestingly, we found that the cell-type with the highest proportion of pre-SWR silence were Martinotti neurons, a subclass of SOM neurons that exert powerful inhibitory control of the pyramidal tuft dendrites in L1 (Mu- rayama et al., 2009; Wang et al., 2004) (Figure 4E,F). Therefore, altogether, lower inhibition and reduced thalamic input to the tuft dendrites appears to be well suited to enhance the selective strengthening of cortical input.

There are a number of technical limitations to note in our study. First, two-photon calcium imaging is an indirect measure of spiking, and importantly, the relationship between calcium signal and firing rate is cell-type dependent (Kerlin et al., 2010). Previous data, and our patch-clamp experiments, indicate that neuronal responses to SWR are not only spatially sparse, but also temporally sparse—that is, a minority of cells fire relatively few action potentials, during a minority of SWR (Csicsvari et al., 1999; Mizuseki and Buzsáki, 2013; Ylinen et al., 1995). Because even the latest calcium indicators struggle to resolve single spikes (Huang et al., 2021), particularly in inhibitory neurons, our data likely underestimates the number of spikes across cell types. Second, the optogenetic silencing of thalamic axons (Figures 6, 7) was likely incomplete. To prevent the red laser light from triggering visual responses by reaching the back of the retina (Nikbakht and Diamond, 2021), we used a low power (2-4 mW/mm^2^) and a small diameter light beam (400 µm). Although optical techniques have limitations in their ability to capture some features of neuronal activity, the cell type-, projection- and compartment-specific nature of viral-genetic tools, as well as the ability to image large neuronal populations, enabled us to capture previously unreported aspects of cortical circuit function during memory consolidation.

A key aspect of our study was that animals were quietly awake, rather than sleeping or anesthetized. Our whole-cell recordings of RSC neurons indicate that SWR-aligned excitation is weak (Figures 1, S1), and unlikely to stand out prominently against the barrage of subcortical excitation that typically accompanies awake behavior and sensory processing. Several prior studies indicate that some of the same circuit mechanisms may be engaged during sleep, but that this is dependent upon SWR coupling with other oscillatory rhythms, such as thalamocortical spindles. In sleeping rats, mPFC units relayed hippocampal inputs preferentially when released from thalamic drive (Peyrache et al., 2011). Yang et al. (2019) reported that SWR-associated silence in the thalamic mediodorsal nucleus was only observed for SWR that are uncoupled to thalamocortical sleep spindles, indicating that thalamic activity may sometimes contribute important information to the cortical memory trace, rather than sensory interference. Cortical neurons are also known to toggle between a more hyperpolarized ‘down’ state and depolarized ‘up’ state during sleep (Constantinople and Bruno, 2011; Neske, 2016; Steriade et al., 2001). SWR are more likely to occur at these transition points (Battaglia, 2004). The patterns of temporal coupling between SWR and other sleep-related oscillations could be indicators of how distinct memory-related processes, such as synaptic strengthening vs weakening, or recall vs consolidation (Joo and Frank, 2018; Roumis and Frank, 2015) are functionally segregated. How cortical circuits are impacted by SWR—both coupled and uncoupled to other oscillations—during sleep, in a cell-type specific manner, has yet to be resolved.

During wakefulness, animals experience periods of locomotion and active engagement with their sensory environment (‘online’ states), punctuated by periods of immobility and disengagement (‘offline’ states). We show that, in the agranular RSC, these shifts are accompanied by changes in cortical circuit activity, such that thalamocortical excitatory input to cortical dendrites and inhibitory neurons decreases dramatically prior to ‘offline’ windows of memory consolidation. We propose that this shift, which takes place 1-2 seconds before hippocampal SWR, represents a functional disconnect of the cortex from subcortical structures—particularly the thalamus. As a result, the cortical circuit prioritizes memory-related excitation to pyramidal den-drites during windows of lowered inhibitory neuron activity. This work has implications for a range of future questions relating to how the thalamus and other subcortical structures interfere versus contribute to the establishment of memory traces in the cortex, and how cortical cell types gate the flow of information.

## ACKNOWLEDGEMENTS

This work was funded by the European Research Council (ERC Starting Grant #639272 to K.V.; Marie Curie IF #753608 to A.R.C.), Research Council of Norway (#231495 to K.V. and #274328 to A.R.C.), and a UiO:LifeScience internationalization grant. We thank Kristin Larsen Sand, Eivind Hennestad, Laura Bojarskaite, Sverre Grødem, Michele Gianatti and Lyle Graham for technical assistance. We thank Bruno Pichler (Independent NeuroScience Services; INSS), for developing the custom two-photon microscope and providing technical assistance. Viral constructs were gifts from Douglas Kim & GENIE project, Hongkui Zeng, and Edward Boyden. Some schematics were created with Biorender.com. Finally, we thank Andrew Alexander, Lukas Fischer, Mark Harnett, Freyja Ólafsdóttir, Aree Witoelar and Jonathan Whitlock for providing critical comments on a draft of the manuscript.

## AUTHOR CONTRIBUTIONS

A.R.C. and K.V. designed research. A.R.C. and C.N.B. performed two-photon imaging experiments. K.V. performed in vitro whole-cell recordings. A.R.C. performed all other experiments and data analysis. A.R.C. and K.V. wrote the paper with feedback from C.N.B.

## Experimental Procedures

### Resource availability

The datasets and code generated in this study are available at [name of repository] and Github, respectively [accession code/web link].

### Mice

Male and female mice between the ages of 8 and 20 weeks were used. For experiments with whole-cell patch clamp recordings, optogenetics, and bulk dendritic imaging, wild-type mice (C57Bl/6J; Janvier labs) were used. For experiments involving cell-type specific two-photon imaging, the following mouse strains were used (with JAX catalog #): Thy1-GCaMP6s (GP4.3 line, #024275) for the targeting of excitatory neurons (Dana et al., 2014), Gad2-IRES-Cre (#010802), VIP-IRES-Cre (#010908), and SOM-IRES-Cre (#018973) for targeting inhibitory neurons (Taniguchi et al., 2011). Surgical procedures were carried out under isoflurane anesthesia (3 % induction, 1 % maintenance), while body temperature was maintained at 37 deg C with a heating pad (Harvard Apparatus). A subcutaneous injection of 0.1 mL Marcaine (bupivacaine 0.25 % *m/V* solution in sterile H2O) was delivered at scalp incision sites. Post-operative injections of analgesic (Temgesic, 0.1 mg/kg) were administered subcutaneously. Mice were housed in a reversed 12 hr light/ 12 hr dark cycle, and experiments were carried out during their dark phase. All procedures were approved by the Norwegian Food Safety Authority (project: FOTS 6590, 7480, 19129). Experiments were performed in accordance with the Norwegian Animal Welfare Act.

### LFP electrode implants and recordings

Mice were first implanted with a titanium headbar. The skin and fascia over the dorsal skull surface were removed, and the headbar was glued to the skull with cyanoacrylic glue and covered with dental acrylic (Jet Denture Repair (Lang Dental) cured with Meliodent Liquid). After recovering from headbar implantation for at least 48 hours, they received LFP electrode implants under isoflurane anesthesia. Burr holes were drilled for the LFP electrode and reference electrode over the dorsal CA1 region of the hippocampus (−2 mm A/P, 2 mm M/L) and contralateral primary somatosensory cortex (−0.5 mm A/P, 3 mm M/L), respectively. Silver wire electrodes (0.125 mm diameter, insulated, GoodFellow) were lowered to a depth of 0.8 mm from the brain surface to target the pyramidal layer of dorsal CA1. The reference electrode was implanted at the brain surface and secured with cyanoacrylate glue. Mice recovered from isoflurane anesthesia while head-fixed for at least 15 minutes, and electrode placement was confirmed by monitoring the LFP signal online with an oscilloscope. Electrodes were secured with cyanoacrylate glue and dental cement.

LFP recordings were band-pass filtered (0.1-1000 Hz) and amplified (1000x) with a DAM50 differential amplifier (WPI Inc). Line noise was removed using a HumBug 50/60 Hz Noise Eliminator (Quest Scientific Inc). For experiments, mice were head-fixed under a two-photon microscope objective after brief (1-2 minutes) isoflurane anesthesia. They were given approximately 20 minutes to recover from anesthesia before recordings were taken. During recordings, mice were able to walk freely on a disc. Experiments were performed in the dark, during the mouse’s dark phase. Data collection and synchronization with image acquisition was carried out using a custom automation program in LabVIEW (National Instruments, using a PCIe-6351X DAQ). LFP and rotary encoder signals were acquired at 20 kHz and downsampled to a final sampling frequency of 2500 Hz.

### SWR detection procedure

SWR were detected based on an automated thresholding of the band-pass filtered LFP (150-250 Hz) (Csicsvari et al., 1999; Mölle et al., 2006; Siapas and Wilson, 1998). We first computed the absolute of the Hilbert transform of the filtered LFP (MATLAB 2018). The output was smoothed by convolving with a 1052-point Gaussian filter with σ = 40 ms. Then, the findpeaks function identified peaks 3 standard deviations above the mean in a moving 1-second time window. A minimum peak width at half height of 15 ms and a minimal peak distance of 25 ms were required.

To avoid the selection of movement-related artifacts, only SWR event candidates co-occurring with a wheel speed of zero were selected for further consideration. Potential SWR waveforms were then visually inspected using the raw LFP trace and the spectrogram. Cross-validation of visual inspection with 2-3 subjects, as well as a recently developed recurrent neural-network based automated tool (Hagen et al., 2021) were used for a subset of recordings.

### In-vitro whole cell patch clamp recordings

Eight-week-old female mice received a stereotactic injection of AAV1-hSyn-ChrimsonR-tdTomato (UPenn Viral Vector Core) to the thalamus (1 x 50 nL; −1.22 mm A/P, 1.25 mm M/L, −2.6 mm D/V). They expressed the virus for at least 5 weeks before they were used for patch clamp experiments.

To optimize slice viability from mature brains, we used the N-Methyl-D-glucamine (NMDG) protective recovery method described in Ting et al. (2018). Briefly, mice were deeply anesthetized with pentobarbital sodium (90 mg/kg, intraperitoneal) and transcardially perfused (10 mL/min) with chilled (2-4 deg C), carbogenated NMDG-HEPES aCSF (in mM): 92 NMDG, 2.5 KCl, 1.25 NaH2PO4, 30 NaHCO3, 20 HEPES, 25 glucose, 2 thiourea, 5 Na-ascorbate, 3 Napyruvate, 0.5 CaCl2·2H2O, and 10 MgSO4·7H2O. The brain was dissected and placed in chilled NMDG-HEPES aCSF for 1 minute, before embedding in agarose.

Coronal slices (300 µm thick) including the RSC (approx. 1.2 mm to 3.2 mm posterior from Bregma) were cut with a vibratome (Leica Biosystems) and transferred to a warm (37 deg C) recovery chamber with NMDG-HEPES aCSF. Na^+^ “spike-in” solution (2M) was introduced at a controlled rate (250 to 1000 µL every 5 minutes for a total of 20 minutes, according to the recommended protocol in Ting et al. (2018) for mice of 3-6 months of age). Finally, slices were incubated in oxygenated artificial cerebrospinal fluid (aCSF; in mM: NaCl 124, KCl 2.5, NaH2PO4 1.2, NaHCO3 24, HEPES 5, Glucose 12.5, MgSO4.7H2O 2, CaCl2.2H2O 2. (308-312 mOsm and pH 7.35) for at least 45 minutes at room temperature before recordings. Recordings were performed at room temperature, and the recording chamber was perfused with oxygenated aCSF. Pipettes were pulled from thin-walled borosilicate capillaries with filament (#BF150-110-10HP; outer diameter 1.5 mm, inner diameter 1.1 mm, Sutter Instruments) using a weighted dual-stage micropipette puller (Narishige PC-10). Pipettes had approximately 2 µm diameter tips and a resistance ranging from 7-9 MOhm when filled with intracellular solution comprised of (g/mol): K Gluconate 234.25, KCl 74.55, HEPES 238.3, MgCl2 95.21, ATP.Mg2+ 507.18, GTP.Na+ 523.18, EGTA 380.35. The pH was adjusted to 7.3 with concentrated potassium hydroxide. During recordings, pipettes were mounted onto motorized micromanipulators (Sensapex).

Neurons were visualized using a custom-build microscope (Thorlabs, building instructions and parts list are available on request) using a 940 nm infrared LED (#M940L3) and oblique illumination (Camera: Thorlabs, Kiralux 2.3). For optogenetic illumination, a 625 nm LED light beam (Thor-Labs, #M625L4) was collimated into the substage condenser lens (Thorlabs, #C330TMD-B) and had a diameter of approximately 3 mm when passing the brain slice (power density was approximately 4 mW/mm^2^). For optogenetic stimulation, we used a 5 ms long pulse after bath application of 4-AP (0.5 M) and TTX (1 mM) to suppress spiking and to enable direct depolarization of the axons (Petreanu et al., 2009). Voltage responses to current pulses from −400 pA to +400 pA were recorded before and after drug application in order to confirm lack of spiking. Signals were amplified (Multiclamp 700B, Molecular Devices) and sampled using a DAQ (National Instruments, PCIe-6351) at 20kHz using WaveSurfer (http://wavesurfer.janelia.org). Series resistance values ranged from 8.5 to 15.6 MOhms. Membrane potentials were not corrected for the liquid junction potential.

### In-vivo whole cell patch clamp recordings

At least 48 hours prior to recording sessions, 8-10 week old female mice were anesthetized with isoflurane and received a titanium headbar implant with a circular well over the dorsal cortex. An LFP recording electrode and reference electrode were also implanted in CA1 and barrel cortex, respectively (see previous section “LFP electrode implants and recordings). The exposed skull surface was then covered with silicone elastomer (Kwik-Cast, WPI) and a thin layer of dental cement. On the day of recording, the dental cement and Kwik-Cast were removed under isoflurane anesthesia, and the exposed skull surface was cooled with a combination of ice and chilled aCSF (see previous section “in-vitro whole cell patch clamp recordings” for aCSF composition). A 1 mm diameter craniotomy was drilled over the agranular RSC while keeping the skull and brain surface cool with continual perfusion of cold saline. The bone flap was gently lifted so as not to damage the underlying dura. The exposed brain and skull surface were covered with agar (1 %, Sigma #A6877), and the mouse was given at least 20 minutes to recover from anesthesia while head-fixed at the whole-cell patching setup. See section “in vitro whole cell patch clamp recordings” for composition of intracellular solution.

During recordings, mice were head-fixed while allowed to walk on a circular disc treadmill. The treadmill could be immobilized during recordings to increase stability. Whole-cell patch-clamp electrodes were introduced into the cortex using a Sensapex micromanipulator, at an angle of approximately 20 deg from perpendicular. For data acquisition we used a DAGAN BVC-700A amplifier and WaveSurfer. The electrode impedance was monitored continuously with 50 % duty cycle, 10 Hz current pulses alternating between 0 and −1.11 nA. For the initial descent, contact with the cortical surface and penetration of the dura, the electrode was advanced with a strong positive pressure of 100-300 mmHg applied to the interior of the pipette, which was quickly reduced to 40-50 mmHg after penetration of the dura. The electrode resistance was then determined by visual inspection and fully compensated by the bridge circuit of the amplifier, and voltage offset adjusted according to an estimated tip offset potential (−14 mV).

The procedure for gaining whole-cell access was taken from Schramm et al. (2014). Briefly, the electrode was advanced in small (5-8 micron) steps until a deflection of the voltage response was observed, indicating contact with a cell. The pressure was quickly released by valve, and a giga-seal formed typically with the aid of light suction applied by mouth to the pipette. The amplitude of the current steps was progressively reduced from −1.11 nA to −110 pA to −10 pA to avoid large voltage changes. When a stable gigaohm-seal was achieved, further light suction was applied to achieve whole-cell access. Once stable whole-cell access was achieved, the amplifier capacitance compensation and access resistance compensation were applied online. Recordings were performed in the dark.

### Viruses, retrograde tracing and cell targeting procedures

For the targeting of inhibitory interneurons, mouse lines were used that expressed Cre recombinase specifically in neurons expressing Gad2, VIP, and SOM (JAX, see “Mice” section for strain information and catalog numbers). In these mice, injections delivering AAV1-CAG-flex-GCaMP6s.WPRE.SV40 (Addgene #100842-AAV1) were made in the agranular (dorsal) region of retrosplenial cortex in the left hemisphere (8-9 x 20 nL injections; RSC coordinates: AP: −2.3 mm; ML: 0.5 mm; DV: −0.2 mm, Drummond Nanoject). Excitatory neurons in all layers were targeted either by using the Thy1-GCaMP6s mouse line (see previous section “Mice” for catalog number; Dana et al. (2014)), or by viral delivery of GCaMP6f under control of the CaMKII promotor (10-12 x 20 nL injections of AAV1-CaMKII-GCaMP6f; UPenn Viral Vector Core).

To target GCaMP6s expression specifically to RSC Layer V pyramidal neurons, in wild-type mice, a single 250 nL stereotactic injection of AAV2-retro-Syn-EBFP-Cre (Addgene #51507-AAVrg; (Madisen et al., 2015; Tervo et al., 2016)) was delivered to the pons in the left hemisphere (AP: −4 mm; ML: 0.4 mm; DV: 5.5 mm), such that cortical Layer V pyramidal tract neurons, one of the two main classes of pyramidal cells, which project to the brainstem and spinal cord, would express Cre recombinase. During the same surgery, 8-9 20 nL injections of AAV1-Syn-flex-GCaMP6s (Addgene #100845-AAV1, Chen et al., 2013) were delivered to the RSC (AP: −2.3 mm; ML: 0.5 mm; DV: −0.2 mm).

For thalamic axon experiments, we first performed retrograde tracing experiments to determine which thalamic nuclei sent projections to agranular RSC. In wild-type mice, a single 100 nL injection of AAV2-retro-GFP (Addgene #37825-AAVrg) was administered to agranular RSC in the left hemisphere (AP: −2.18 mm; ML: 0.5 mm; DV: −0.2 mm), and the mice expressed the virus for 4-5 weeks before perfusion and sectioning for histology. To enable imaging of neural activity from thalamic projections, AAV1-Syn-GCaMP6s-WPRE-SV40 (Addgene #100843-AAV1; Chen et al., 2013); was delivered to the thalamus (AP: −1.22 mm; ML: 1.1 mm; DV: −2.6 mm) via a single 50 nL stereotactic injection, and axons in L1 of agranular RSC (up to 80 µm depth from pial surface) were subsequently imaged. For in vitro or in vivo optogenetic manipulation of thalamic projections, single 50 nL stereotactic injections of either AA1-Syn-ChrimsonR-tdTomato (Addgene #59171-AAV1; (Klapoetke et al., 2014)) or AAV5-Syn-GFP-Jaws (Addgene #65014-AAV5; (Chuong et al., 2014)) were delivered to the same coordinates.

### Two-photon calcium imaging

Several weeks prior to imaging, mice underwent surgery to receive headbar and glass cranial window implants, as well as virus injections, if necessary. A circular craniotomy approximately 2.5 mm in diameter was made over the left and right agranular RSC (centered on the midline, approx. 2.2 mm to 2.5 mm posterior to Bregma), to allow imaging from a dorsal approach. The skull surrounding the craniotomy (approx. 0.5 mm from craniotomy edge) was thinned to accommodate a glass window implanted in the craniotomy. The window consisted of a circular outer window (3.5 mm diameter, #1.5 coverslip glass) affixed to a half-circle inner window (2.5 mm diameter, #1.5 coverslip glass) with optical adhesive (Norland Optical Adhesive; ThorLabs NOA61), designed to fit over the left RSC, against the central sinus, to prevent dural growth. The window was held in place with gentle pressure so that the outer window fit into the thinned skull area and was flush with the surface of the skull. Heated agar (1 %; Sigma #A6877) was applied to seal any open spaces between the skull and edges of the glass window, and the window was affixed to the skull with cyanoacrylate glue.

For experiments involving imaging across layers, a glass microprism (1×1 mm prism face, Edmund Optics, N-BK7 #86-621; Andermann et al., 2013) was inserted into the left RSC, facing the anterior direction. After the craniotomy was performed, a small slit was made in the dura with a dissecting knife, into which the prism, glued to the underside of the circular cranial window with optical adhesive, was inserted to approximately 1.1 mm depth. After implantation, the mice were given at least 5 weeks to recover before imaging experiments were performed. Images were taken at least 100 µm anterior to the prism face.

Imaging experiments were performed using a custom-built two-photon microscope (Independent NeuroScience Services) with a resonant scanner. The light source was a MaiTai DeepSee ePH DS (Spectra-Physics), and the objective was a 16x water immersion lens with 0.8 NA (Nikon). Images were acquired at 31 Hz using SciScan software implemented in LabView (National Instruments). Laser power was 20-100 mW at the brain surface.

For somata, regions-of-interest (ROIs) were hand-drawn around GCaMP6-expressing neurons whose nuclei were not filled with fluorophore, and signals extracted using custom code in MATLAB (MathWorks, Inc). For ROIs drawn around neuronal somata, signals were adjusted for neuropil contamination by subtracting the median fluorescence of pixels surrounding the ROI (within a distance surrounding the ROI equivalent to the ROI radius) that were not marked as part of another neighboring ROI. Neural activity, defined as *f* (*t*) = Δ*F* (*t*)*/F*_0_, was calculated for all ROIs by first estimating baseline fluorescence (*F*_0_) as the 20^th^ percentile of the activity vector for each ROI. For bulk dendritic recordings, activity was averaged over the entire field of view. For neuronal somata, axons, and dendrite branches, deconvolution of calcium events was carried out as described in Pnevmatikakis et al. (2016), using adaptive time constants in order to capture heterogeneity across neurons in a field of view. To exclude small values likely representing noise, all deconvolved DF/F values under 10^−8^ were set to zero. Then, k-means clustering was used to separate the remaining deconvolved DF/F values of each ROI into two clusters (signal and noise). A linearly spaced vector was calculated between the centroids of the clusters. A threshold was set at the 15^th^ percentile of this vector, and all values under this threshold were set to zero.

### Imaging of calcium-independent fluorescence changes

To measure changes in fluorescence in two-photon imaging experiments that are not due to neural spiking activity, we performed control imaging sessions using an excitation wavelength of 810 nm, based on the isosbestic point of GCaMP (Chen et al., 2013) (approx. 410 nm). For each control session, the laser intensity at both 810 nm and 950 nm was set such that the mean fluorescence intensity over the FOV in both cases was matched. Corresponding DF/F responses were calculated for 810 nm sessions and compared to the calcium-dependent sessions for the same FOV.

### Auto-segmentation procedure for axonal imaging analysis

A custom, automated procedure for ROI drawing (MATLAB 2018b, see previous section “Resource availability” for Github repository) was implemented to select axonal segments in images of thalamocortical projections. The image stack was downsampled and smoothed in time, followed by background subtraction and thresholding. Thresholded components were selected across frames and merged if they spatially overlapped, beginning with the overlapping segments that appeared most often and ending with those that appeared most rarely. A maximum of 300 components were selected from each field of view. Those that fell within 15 pixels of the image boundary were excluded.

To avoid multiple sampling of the same axon branches, a clustered correlation matrix was created from all session-wide response vectors from all autosegmented ROIs (number of clusters = 10). Those that fell within clusters with the highest correlation values in the session (at least 1 standard deviation above the overall mean correlation value) were merged and treated as a single ROI.

### In vivo optogenetics and analysis of optogenetic modulation

Male and female WT mice received single 50 nL injections of either AAV1-hSyn-ChrimsonR-tdTomato or AAV5-hSyn-GFP-Jaws (see previous section “Viruses, retrograde tracing, and cell targeting procedures”) to the thalamus in one hemisphere (AP: −1.22 mm; ML: 1.1 mm; DV: −2.6 mm). Optogenetic stimulation in vivo was performed through the glass cranial window implanted over RSC ipsilateral to the thalamic injection site. 625 nm laser light pulses were delivered through the two-photon microscope objective during simultaneous optogenetic/two-photon stimulation and imaging sessions. To implement this, we mounted a red laser diode (Oclaro, 638 nm, 700 mW) in the position where normally the red channel photomultiplier would be located. To make the beam shape more circular, we focused the light into an optical fiber (Thorlabs, #M15L01) using a fiber coupler with collimation lens (Thorlabs, #PAF2S-11B). The beam was then focused on the objective back aperture such that it diverges when entering the brain. The diameter of the beam was approximately 400 µm (close to the size of the imaging FOV). Laser power of this beam was measured prior to each experiment and was kept in the range of 3-4 mW/mm^2^. To prevent the intense red light from entering the photomultiplier of the green channel, we mounted two OD 6 filters (#ET510/80, Chroma) in series after the secondary dichroic (#T565lpxr, Chroma). For experiments involving Jaws, the optogenetic pulse was a 3-second continuous light pulse. For Chrimson-R, a 5 Hz, 2.5 % duty cycle sinusoidal stimulus (3 seconds in total duration) with 100 ms on/off ramps was used. Each session of combined imaging and optogenetics consisted of 35 trials over 10 minutes, with a varying intertrial interval ranging from 8 to 15 seconds. Experiments were performed with a red masking light positioned approximately 10 cm in front of the mouse’s face. In order to test for significant effects of laser stimulation, the average response magnitude in the middle 1/3 of the light stimulus time period (+1 s to +2 s from laser onset) was subtracted from the average response in the pre-laser stimulus period (−2 s to −1 s from laser onset), and the distribution of light stimulus responses was tested for significant mean deviation from zero using a one-sample *t*-test.

### Histology

Mice were deeply anesthetized with pentobarbital sodium (90 mg/kg) and, once the absence of reflexes was detected, transcardially perfused with 4 % paraformaldehyde (w/v) in phosphate buffered saline (PBS). Brains were kept for at least 24 hours in 4 % paraformaldehyde, then cryoprotected with 30 % sucrose (w/v) in PBS solution for 24-48 hours. Brains were then embedded, frozen and cut into 30 µm coronal sections with a cryostat. Sections were DAPI stained prior to mounting.

### Analysis of significant modulation

All sessions with fewer than 20 SWR remaining after the locomotion control (see section “Locomotion analysis”) was applied were excluded from analysis. SWR occasionally appeared multiple times within hundreds of milliseconds of each other. Therefore, to avoid confounding effects caused by analyzing SWR clusters, if SWR occurred with less than 2 seconds in between, only the first SWR was included in the analysis.

For analysis of subthreshold Vm modulations in whole cell recordings, spikes were first removed using a combination of a lowpass filter (300 Hz) and a median filter (20 ms). The mean Vm response from −3 to +3 seconds from SWR peak was calculated for each cell. Then, we calculated the upper and lower 95 % confidence interval of the Vm response in the time window −3 to −2 seconds prior to SWR peak. If the mean SWR response for a cell exceeded this interval within a time window of 500 ms beginning at SWR peak, the cell was considered as having a synaptic SWR response.

To investigate slow, pre-SWR modulations in Vm, spiking activity and deconvolved DF/F, for individual cells or subcellular components (e.g. dendrite branches or axon segments) the average of each SWR-by-SWR response matrix was taken between −2 s and +500 ms from SWR peak, and compared to a shuffled version (adapted from Jadhav et al. (2016)). Each trial (individual SWR response) was circularly shifted by a random amount of up to 2 seconds. Responses to the same SWR were shifted by the same random fixed amount, thus preserving the temporal structure of the response but shifting it relative to SWR. The mean squared error of this shuffled matrix was calculated, then compared to the mean squared error of the actual SWR response matrix. If the mean squared error was smaller than that between the actual response matrix and the mean shuffled matrix, 95 % of the time across 1000 instances of shuffling, the cell was considered modulated. Suppression or activation was determined by comparing the mean response in a baseline period (−3 s to −2 s from SWR) versus a test period (−1 s to 0 s from SWR), for all significantly modulated cells.

In addition to investigating SWR modulations on an individual cell basis, we also analyzed the size of SWR modulations across populations of neurons or subcellular components. To analyze the size of pre-SWR modulations (except for bulk dendritic DF/F), we calculated the mean difference of the response from −3 to −2 seconds from SWR (baseline period) and the response from −1 to 0 seconds from SWR (pre-SWR period). For SWR modulations, we calculated the mean difference of the response from −1 to 0 seconds from SWR (pre-SWR period) and the response from 0 to +500 ms from SWR (SWR period). For bulk dendritic imaging, we analyzed DF/F, which has a poorer temporal resolution compared to spiking activity or deconvolved DF/F. Therefore, to analyze bulk DF/F response to SWR, we compared activity at SWR peak (0 to +100 ms from SWR) to a 500 ms time window afterwards (+150 ms to +650 ms from SWR).

To test significance across populations of cells, we assumed normality and used one-sample *t*-tests, except in the case of whole-cell recordings with small sample sizes (<30 cells, Figure 1), where a non-parametric test (one-sample Wilcoxon signed rank test) was used. We used the Bonferroni method to correct for multiple comparisons.

### Locomotion analysis

To restrict our analysis to include only those SWR that occurred during prolonged periods of immobility, we subdivided each 10-minute experimental session into locomotion and immobility epochs (Figure S2). Locomotion was defined as an instantaneous wheel speed of > 1 cm/s. Bouts of immobility lasting less than 2 seconds, as well as a buffer period of 2 seconds before and after each locomotion bout, were categorized as locomotion. SWR were only included in the analysis if they did not occur within 3 seconds of a locomotion bout.

## Supplementary Figures

**Figure S1.**
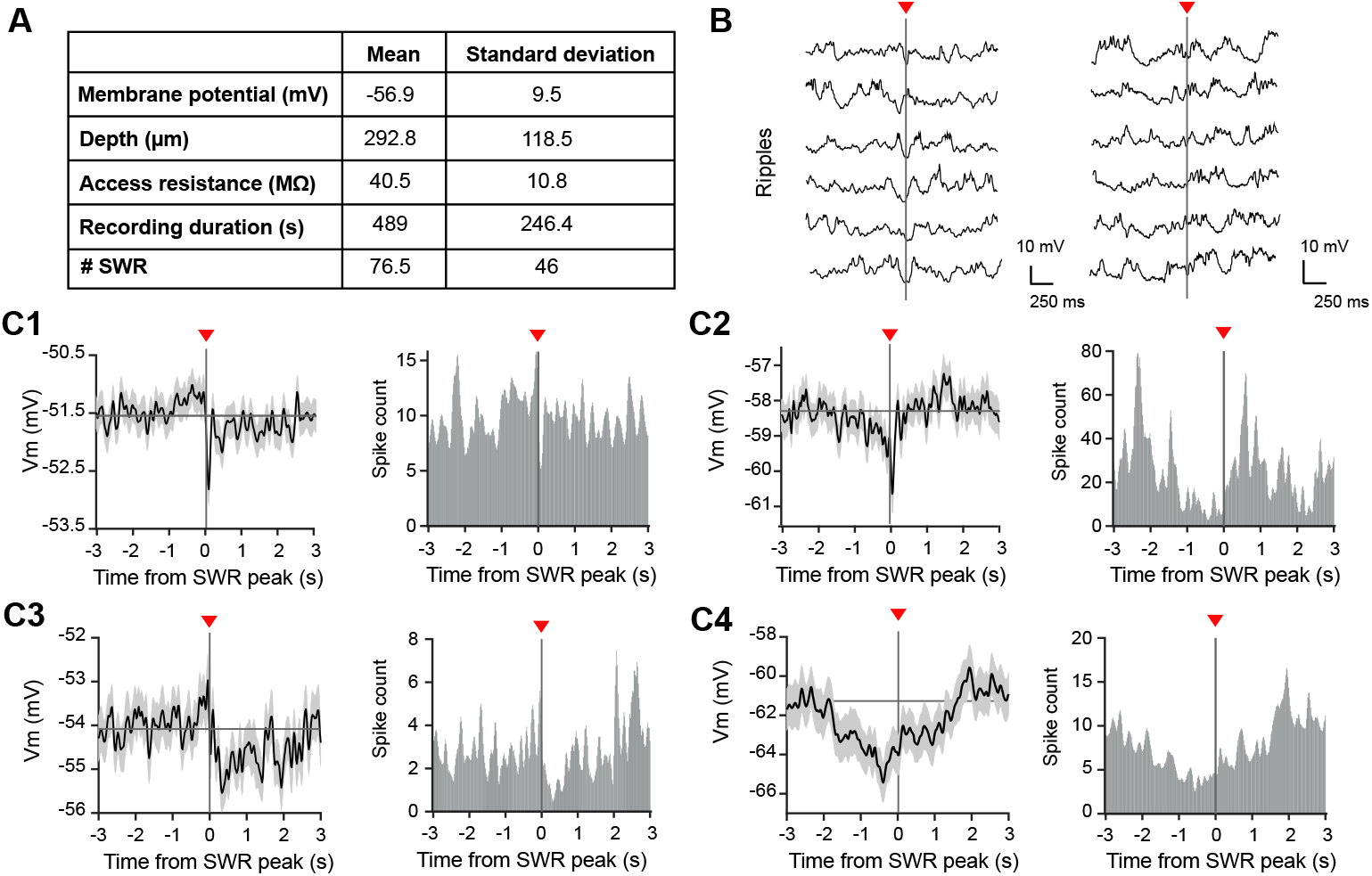
SWR-associated subthreshold Vm modulations are not always reflected in spiking output. (**A**) Table of information on whole-cell recordings (30 cells, 11 mice). (**B**) Example Vm traces from two cells (left and right), 1 sec before and after SWR (6 instances for each cell). Spikes have been removed (see Experimental Procedures). Red arrow indicates timing of SWR peak. Example cell on the left is the same as in C1, cell on the right is the same as C4. (**C1-4**) Average ± s.e.m. subthreshold Vm (left) and spike histogram (right) for 4 example cells. Red arrow indicates timing of SWR peak. Gray horizontal line indicates Vm baseline. Bin size = 20ms.

**Figure S2.**
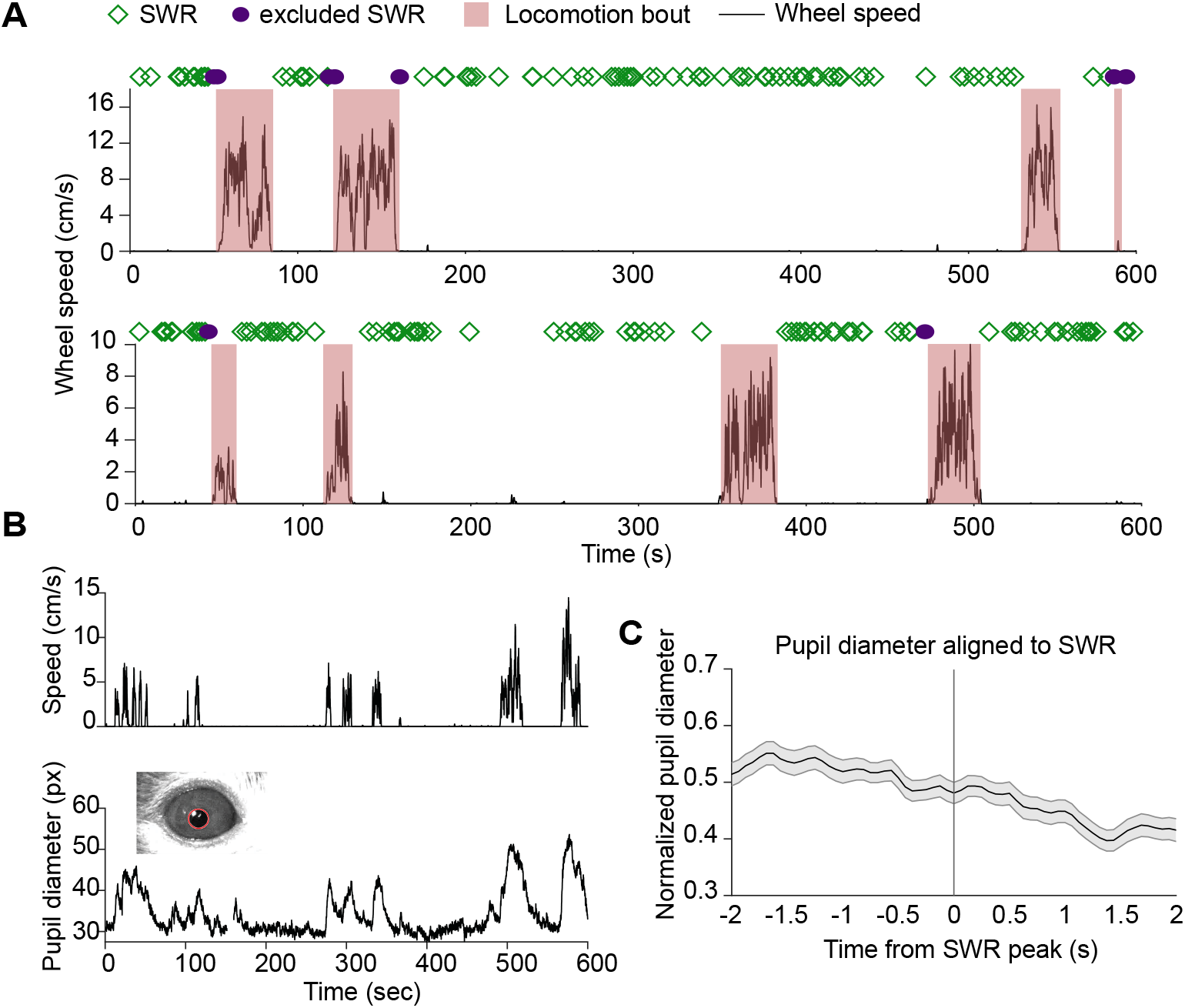
Wheel speed and pupil diameter as readouts of locomotion and arousal during two-photon imaging. (**A**) Typical examples of locomotion speed during a 10-minute imaging session from two different mice. Bouts of locomotion are labeled with red boxes and SWR occurrences are marked with green diamonds. SWR that were removed from the analysis due to their proximity to a bout of locomotion are labeled in purple. 99 SWR (top); 77 SWR (bottom). (**B**) Locomotion speed (top) and pupil diameter (bottom, units = pixles) during a 10-minute imaging session. Inset shows the pupil tracking. (**C**) Average ± s.e.m. pupil diameter (pixels), normalized and aligned to SWR peak (152 SWR, 2 mice).

**Figure S3.**
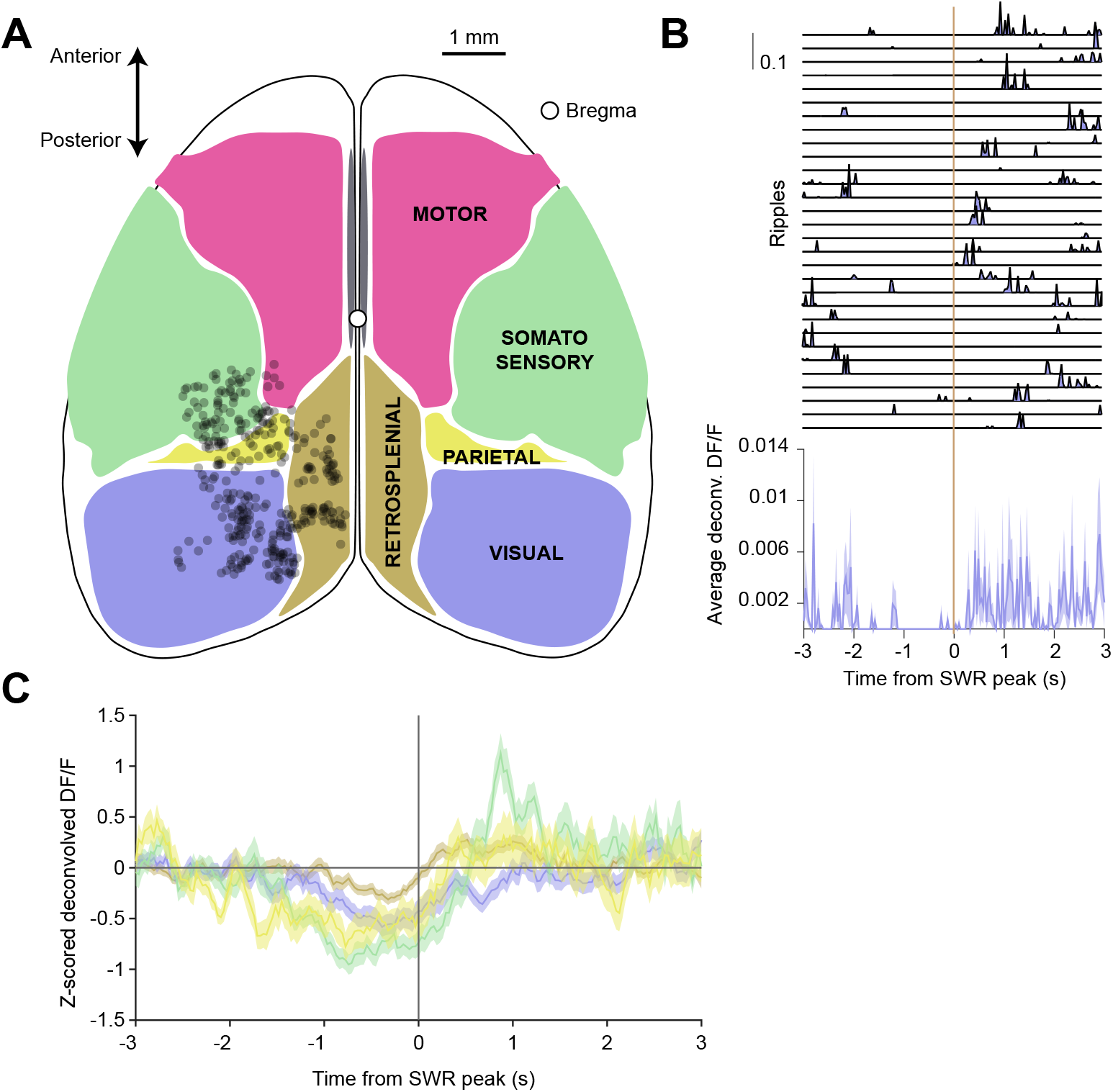
Pre-SWR reductions in activity occur in L1 inhibitory neurons across several neocortical regions. (**A**) Dorsal view of cortex with major sub-divisions according to Kircaldie et al. (2012) (5 mice). Black circles denote the locations of L1 inhibitory neurons showing significant pre-SWR reductions in activity (218 cells). (**B**) Deconvolved DF/F responses, across SWR events (top), and averaged SWR-aligned (bottom), of an example L1 inhibitory neuron in visual cortex. (**C**) Averaged ± s.e.m. deconvolved DF/F for all cells in (A) separated by cortical area. Colors correspond to the regions in (A).

